# Neural Signatures of Evidence Accumulation Encode Subjective Perceptual Confidence

**DOI:** 10.1101/2023.04.28.538782

**Authors:** Wei Dou, Lleymi J. Martinez Arango, Olenka Graham Castaneda, Leopoldo Arellano, Emily Mcintyre, Claire Yballa, Jason Samaha

## Abstract

Confidence is an adaptive computation when environmental feedback is absent, yet there is little consensus regarding how perceptual confidence is computed in the brain. Difficulty arises because confidence correlates with other factors such as accuracy, response time (RT), or evidence quality. We investigated whether neural signatures of evidence accumulation during a perceptual choice predict subjective confidence independently of these factors. Using motion stimuli, a central-parietal EEG component (CPP) behaves as an accumulating decision variable that predicts evidence quality, RT, accuracy, and confidence (Experiment 1). Psychophysically varying confidence while holding accuracy constant (Experiment 2), the CPP still predicts confidence. Statistically controlling for RT, accuracy, and evidence quality (Experiment 3), the CPP still explains unique variance in confidence. The results indicate that evidence accumulation, indexed by the CPP, is tightly linked to the subjective perceptual experience of sensory information. Independent of other factors, pre-decision neural signatures of evidence accumulation encode subjective confidence.

## Introduction

When humans make decisions they are typically able to introspect about the probability that a choice was correct and provide a subjective judgment of their own confidence. In the absence of external feedback, confidence judgments are thought to serve an important functional role in adaptive behavior. Insofar as confidence reflects uncertainty in one’s choice, it can help guide future decision policies (Desender et al., 2018; Rollwage et al., 2020; Talluri et al., 2018; van den Berg et al., 2016) or be used to adaptively weight the integration of different information sources into a single choice (Braun et al., 2018; Samaha et al., 2019; Urai et al., 2019). For instance, one might decide to allocate resources to collecting more evidence when they have low confidence in their initial choice (Desender et al., 2018; Kepecs et al., 2008; Yeung & Summerfield, 2012). Moreover, confidence has played an important role in theorizing about conscious perception with some arguing that perceptual awareness involves a meta-cognitive component akin to introspection (Brown et al., 2019; Y. Ko & Lau, 2012) and that confidence judgments can sometimes signal the presence of conscious perception better than the ability to make correct perceptual choices (Michel, 2022).

A growth in interest in confidence in recent decades has given rise to many different models of the underlying computations (Aitchison et al., 2015; Fetsch et al., 2014; Fleming & Daw, 2017; Kepecs et al., 2008; Kiani et al., 2014; Kiani & Shadlen, 2009; Maniscalco & Lau, 2016; Meyniel et al., 2015; M. A. K. Peters, 2022; Pleskac & Busemeyer, 2010; Pouget et al., 2016; Sanders et al., 2016). There is currently little consensus on which models best capture human confidence behavior (Rahnev et al., 2022) which has given rise to a wide variety of proposals regarding how confidence is encoded in neural activity. Some have argued that confidence is read out in an approximately Bayesian fashion from population-level representations of uncertainty within sensory areas, which naturally explains the close empirical link between confidence judgments and decision accuracy (Geurts et al., 2022; Khalvati et al., 2021; Meyniel et al., 2015). Yet other studies have approached the question from an evidence accumulation perspective, highlighting the contributions of decision-related brain dynamics to confidence (Desender, Donner, et al., 2021; Fetsch et al., 2014; Gherman & Philiastides, 2015; Kiani et al., 2014; Kiani & Shadlen, 2009). According to many evidence accumulation models, confidence can be read out from not only the amount of evidence accumulated for a particular choice but also by the time it took for that much evidence to be acquired, thus linking confidence to accuracy as well as response times (Fetsch et al., 2014; Kiani et al., 2014; Zylberberg et al., 2016). Some evidence accumulation models have also highlighted the possibility that confidence is not based on the exact same evidence that underlies one’s choice, but rather on evidence that continued to accumulate after one’s choice, illuminating the means by which a decision-maker may change their mind (Desender et al., 2021; Desender et al., 2021; Navajas et al., 2016) .

One difficulty in pinpointing neural correlates of confidence is that confidence judgments are typically tightly coupled to the accuracy of one’s decision and to the quality of perceptual evidence, such that easier trials or trials where observers acquired better evidence are accompanied by higher levels of confidence. This means that putative signatures of confidence in the brain could potentially reflect task difficulty or signal quality rather than subjective confidence per se (Lau & Passingham, 2006; Peters et al., 2016; Peters et al., 2017; Samaha, 2015). However, recent psychophysical work has uncovered systematic deviations between confidence and accuracy that can be exploited in order to experimentally manipulate confidence separately from task performance (Koizumi et al., 2015; Maniscalco et al., 2016; Samaha et al., 2016, 2019; Samaha & Denison, 2022; Zylberberg et al., 2012). For example, when confidence and accuracy were dissociated, neurons in the superior colliculus that were previously thought to encode confidence in an oculomotor decision were found to primarily track decision difficulty, but not the animal’s confidence behavior (Odegaard et al., 2018).

Here, we studied neural correlates of perceptual confidence in humans across three experiments combining recordings of electrical brain activity (EEG) and psychophysical manipulations of confidence, accuracy, reaction time, and the quality of perceptual evidence. We focused on an event-related potential (ERP) component termed the central parietal positivity (CPP) which has previously been found to encode the (unsigned) accumulated evidence for a perceptual decision (Kelly et al., 2021; Kelly & O’Connell, 2013; O’Connell et al., 2012, 2018). In particular, the build-up rate (or slope) of the CPP has previously been found to correlate with evidence strength, reaction time, and accuracy and is thought to reflect an amodal decision signal in that it shows similar dynamics and has the same parietal scalp distribution regardless of the sensory modality or motor response demands (O’Connell et al., 2012). The CPP has also been shown to correlate with confidence judgments (Gherman & Philiastides, 2015; Herding et al., 2019a; Vafaei Shooshtari et al., 2019) although not yet when confidence is experimentally dissociated from accuracy. Across three different experiments we find CPP slope, prior to choice initiation, to be a reliable predictor of subjective perceptual confidence, bolstering the notion that humans map the pre-decision evidence accumulated in a given period of time onto a subjective assessment of confidence. However, they do so in such a way that allows confidence to be divorced from the actual accuracy of one’s decision, in violation of some normative Bayesian proposals (Geurts et al., 2022; Meyniel et al., 2015; Sanders et al., 2016).

## Results

### A neural signature of perceptual evidence accumulation

We first sought to identify a neural signature of perceptual evidence accumulation by manipulating the quality of stimulus evidence and testing the CPP for three features observed in prior work (Kelly & O’Connell, 2013; O’Connell et al., 2012). Namely, if the CPP reflects (unsigned) evidence accumulation, its slope (or build-up rate) should increase with the quality of stimulus evidence, should decrease with increasing response times (RT), and should increase for correct compared to incorrect choices. We then sought to test whether CPP also predicted confidence ratings provided for each decision. To this end, we adopted a random dot motion (RDM) task in which a field of moving dots were presented inside a circular aperture, with a fraction of dots moving coherently to the left or right and the rest moving randomly (Figure. 1A). In Experiment 1 we manipulated the strength of sensory evidence by varying motion coherence between 1%, 4.5%, 8%, 12%, 25%, or 40% randomly across trials, whereby higher coherence indicates a stronger motion signal in one direction or the other. After a stimulus presentation (300 ms), participants (*n* = 25) were asked to simultaneously report the global direction of motion (left or right) and their confidence (1-4 scale) using a single key press as quickly and accurately as possible. As shown in Figure 1B, increasing motion coherence led to higher confidence (*F*(5,120) = 91.32, *p <* .001), faster response times (*F*(5,120) = 74.45, *p* < .001), and higher accuracy (*F*(5,120) = 388.16, *p* < .001), consistent with evidence accumulation model predictions of the RDM task as described in prior work (Fetsch et al., 2014).

**Figure 1.**
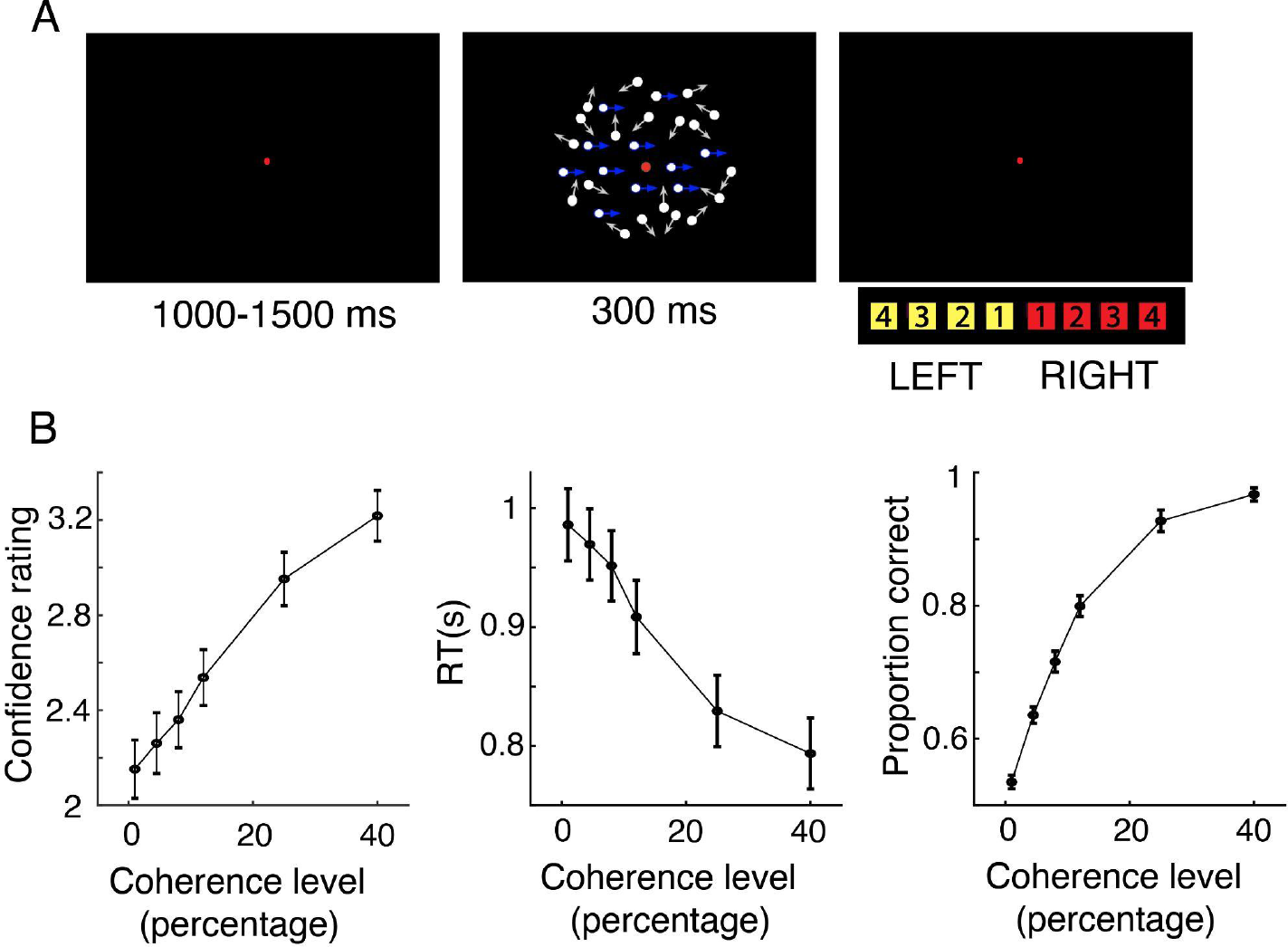
Example trial and behavioral results for Experiment 1 (*n* = 25). (A) Each trial commenced with a central red fixation dot for 1000 to 1500 ms. A random dot motion (RDM) stimulus was then presented for 300 ms (motion coherence was sampled randomly on each trial from the following values: 1%, 4.5%, 8%, 12%, 25%, or 40%) in either a leftward or rightward direction. Participants reported the motion direction (left vs. right) and their confidence (1-4 scale) in the decision simultaneously with a single button press. (B) confidence rating (left) and proportion correct (right) increased with higher coherence level and RT (middle) decreased. Error bars denote ±1 SEM (across subjects). Arrows in panel A represent example dot motion trajectories and were not shown in the actual displays.

During decision formation, we observed a positive ERP component over central parietal electrodes whose activity peaked around 500 ms post-stimulus (Figure 2, *left*) and ramped up prior to the motor response (Figure 2, *right*), resembling the CPP (O’Connell et al., 2012; Twomey et al., 2015). If this signal encodes accumulated evidence, then the relevant parameter for capturing the rate of evidence accumulation is the slope of the CPP component. To this end, we fit a line to each participant’s average CPP waveform using a 200 ms sliding window advanced in 10 ms steps between 100 ms to 550 ms relative to the stimulus-aligned CPP waveform and -1000 ms to -10 ms relative to the response-aligned CPP waveform. We then fit separate linear models predicting CPP slopes at each time window by motion coherence level, RT (divided into 5 quantiles), accuracy (correct or incorrect), and confidence (high or low based on a participant-specific mean-split). For each model we compared the regression slope at each time window to zero (one-tailed) using a nonparametric, cluster-based permutation test to correct for multiple comparisons across time (See “Methods”; Maris & Oostenveld, 2007).

**Figure 2.**
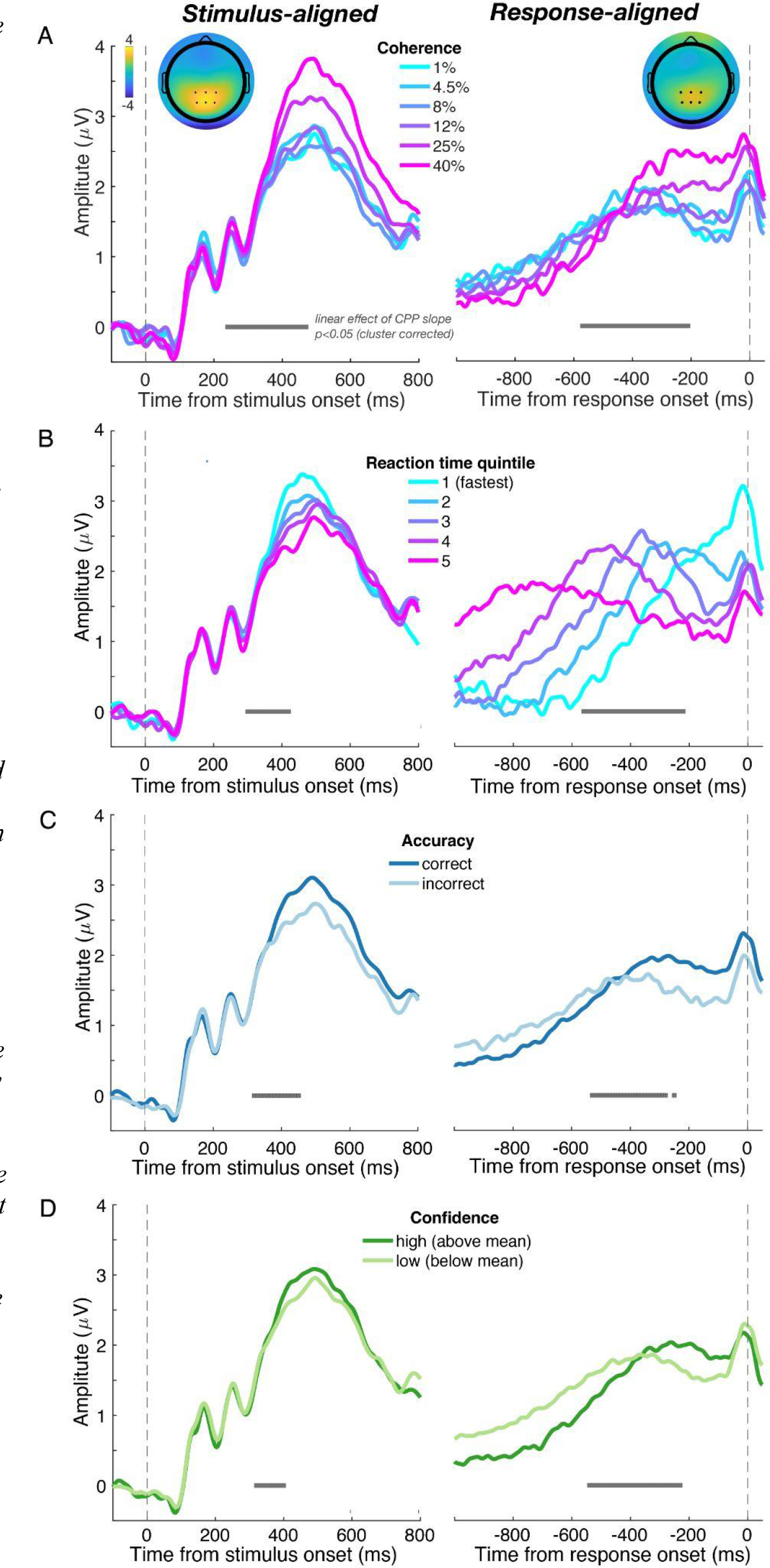
Grand average ERP time-courses aligned to both the target onset (left column) and response execution (right column) for Experiment 1 (*n* = 25). The vertical dashed lines denote the time of target onset (left column) and the time of response execution (right column). (A) the inset topography of the ERP measured after target onset (left; 250 ms to 450 ms) and prior to response execution (right; -510 ms to -130 ms) shows a positive central-parietal component (the CPP) highlighted by the black dots, which denote electrodes used in all subsequent analysis. Both stimulus-aligned and response-aligned CPP slopes increased as motion coherence level increased. (B) Both stimulus-aligned and response-aligned CPP exhibited a slope that scaled with RT (5 equally-sized RT bins). (C) Both stimulus-aligned and response-aligned CPP slopes were higher on correct trials than on incorrect trials. (D) Both stimulus-aligned and response-aligned CPP slopes were steeper on high confidence (above mean) trials than on low confidence (below mean) trials. Gray points below each ERP denote the center of 200 ms time windows in which a linear effect of CPP slope as a function of coherence, RT, accuracy, or confidence reached significance (*p* < .05) after correcting for multiple comparisons across time.

As shown in Figure 2A, this analysis showed a significant positive cluster spanning 240 to 470 ms post-stimulus and a significant cluster spinning -570 to -210 ms prior to response execution whereby steeper CPP slopes were associated with higher coherence levels. As shown in Figure 2B, CPP slopes were significantly negatively correlated with RT bin within a cluster from 300 to 420 ms for the stimulus-aligned CPP and a cluster from -560 to -220 ms for the response-aligned CPP, indicating that the CPP ramps up quicker for faster RTs. Figure 2C shows that CPP build-up rate was also significantly higher for trials where participants correctly discriminated the direction of motion compared to incorrect trials for a cluster spanning 320 to 450 ms post-stimulus onset and a cluster spinning -530 to -280 ms prior response onset. Lastly, Figure 2D confirms the prediction that CPP slope was also related to confidence judgments. Compared to low confidence trials, CPP ramped up significantly faster on high confidence trials during a cluster spanning 320 to 400 ms post-stimulus and from -540 to -230 ms prior to the response.

One interpretation of these results is that subjective confidence judgments can be read out from the neural signature of perceptual evidence accumulation prior to the response. However, in Experiment 1, high confidence trials were also associated with stronger evidence quality and higher accuracy. Thus the relationship between CPP slope and confidence might be confounded by decision accuracy. We therefore sought to tease apart the contributions of task performance and confidence to the CPP build-up rate in Experiment 2.

### Isolating neural signals of perceptual confidence from decision accuracy

We aimed to experimentally manipulate subjective confidence independently of task accuracy using the so-called “positive evidence bias” (PEB) phenomenon that has been observed repeatedly in human (Desender et al., 2018; Ko et al., 2022; Koizumi et al., 2015; Maniscalco et al., 2016; Rollwage et al., 2020; Samaha et al., 2016, 2016, 2019; Zylberberg et al., 2012), monkey (Odegaard et al., 2018), and rodent (Stolyarova et al., 2019) confidence behaviors. The PEB refers to the fact that decision-makers tend to overweight the overall magnitude of evidence in favor of a choice when rating confidence, rather than rely on the actual balance of evidence for both choice alternatives. In Experiment 2 (*n* = 25) we therefore designed the RDM stimulus so that it contained two levels of “positive evidence” or PE (i.e., motion signal supporting the correct choice) and with individually-adjusted levels of “negative evidence” or NE (motion signal supporting the incorrect choice) such that accuracy was matched across the two conditions. Concretely, we set the motion coherence for the correct choice direction to be 50% in the “high PE” condition and 25% in the “low PE” condition. We then found the level of NE for each subject and PE level such that accuracy (% correct) approximated 79% in both conditions using a 1-up, 3-down adaptive staircase prior to the main task (see Methods; Figure 3A).

**Figure 3.**
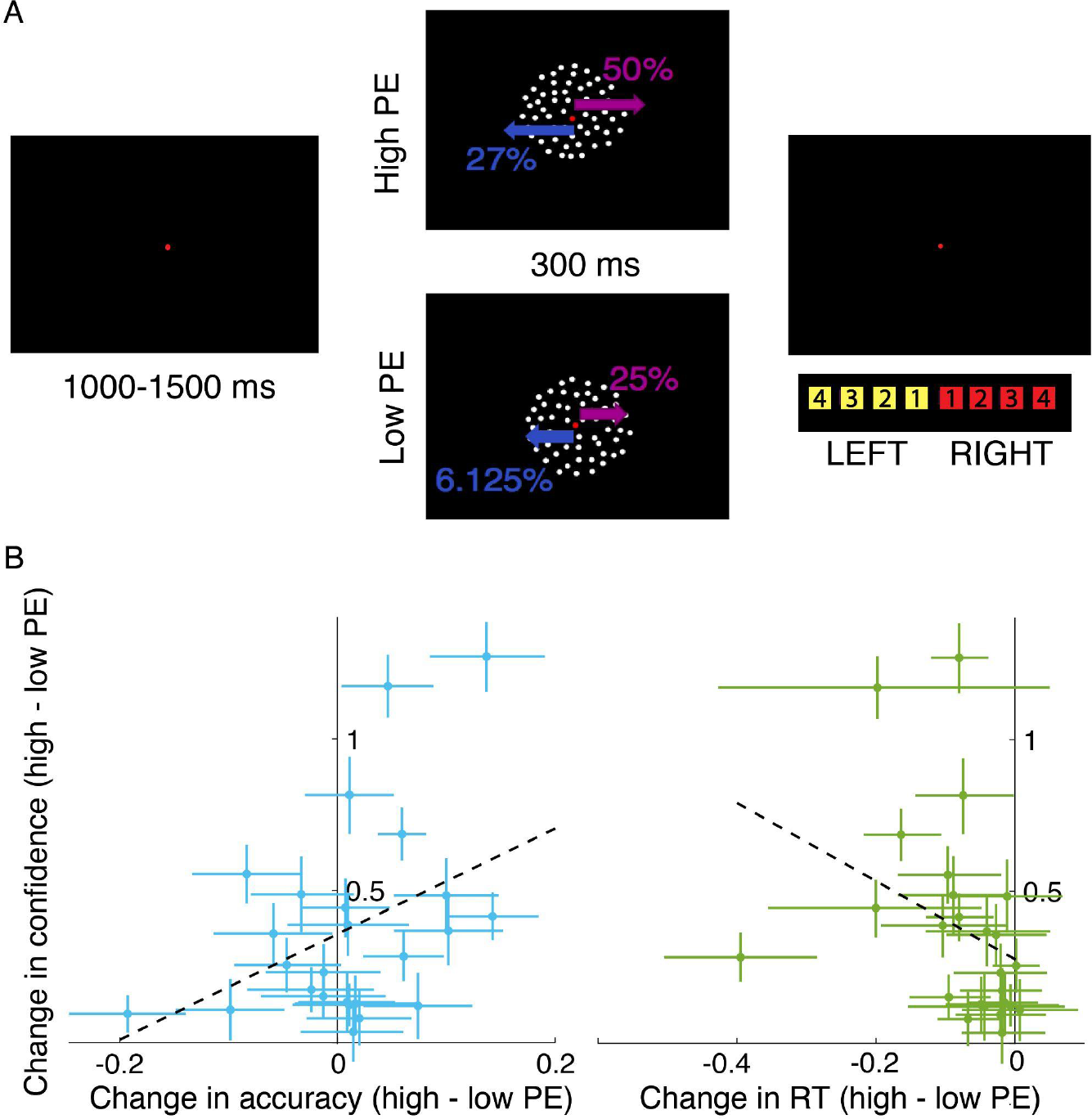
Example of an experimental trial and behavioral results for Experiment 2 (*n* = 25). (A) Following fixation, a RDM stimulus which contained either high or low “positive evidence” (PE) was shown for 300 ms. High PE stimuli always contained a 50% coherent motion signal moving in the correct direction whereas low PE stimuli contained 25% coherent motion moving in the correct direction. “Negative evidence” levels (NE; or motion opposing the correct direction) were varied for each participant and PE level separately to achieve ∼79% discrimination accuracy (in this example participant 25% NE was required in the high PE condition and 6.25% NE was required in the low PE condition). Participants then reported the motion direction (left vs. right) and their confidence (1-4 scale) in the decision simultaneously with a single button press. (B) The effect of PE on confidence (high - low PE; y-axis) and the effect of PE on the proportion of correct responses (high - low PE; x-axis) is shown on the left plot. The same for RT is shown on the right plot. Dots represent individual participants and error bars are bootstrapped 95% confidence intervals. The dashed lines represent the least-squares fit. The left plot shows that participants were approximately equally accurate for high and low PE stimuli, yet all participants showed higher confidence following high compared to low PE stimuli, demonstrating a dissociation between confidence and accuracy. Moreover, the PE effect on confidence was not significantly correlated with the PE effect on accuracy across participants (*rho* = .27, *p* = .19). The right plot shows that the majority of the participants were faster for high PE than for low PE and this difference was correlated negatively with the PE effect on confidence (*rho* = -.55, *p* = 0.005), suggesting confidence was not dissociated from RT in this experiment.

We anticipated that, during the main task, accuracy would be approximately equated for high and low PE, but, if the PEB for confidence holds for our stimuli, then confidence would be higher in the high PE condition. Figure 3B shows the difference in accuracy between high and low PE for each observer on the x-axis (with boostrapped 95% confidence intervals) and the same difference (high - low PE) in confidence on the y-axis. The staircase was successful in controlling accuracy across the two PE conditions at the group level. Low PE accuracy was, on average (±SEM) 77±1.5% and high PE accuracy was 78±2.0%. As can be seen in Figure 3B, a few participants were more accurate for high PE and a few were less accurate, but the majority overlapped zero such that, at the group-level accuray did not significantly differ across PE levels (*t*(24) = -0.69, *p* = 0.50). In contrast, every participant showed numerically higher confidence following high (mean confidence = 3.0±0.11) compared to low PE (mean confidence = 2.6±0.12) stimuli with 23/25 participants showing a significant (*p* < 0.05) elevation in confidence (Figure 3B; bootstrap test), leading to a large effect at the group level (*t*(24) = -5.77, *p* < .001). If confidence were merely a more sensitive estimator of task difficulty, then we may expect the PE effect on confidence to predict the PE effect on accuracy across participants, and although there was a small positive correlation, this was not near statistical significance (*rho* = .27, *p* = .19). This indicates that it is not necessarily the case that an individual who happened to have higher accuracy on high PE trials also had higher confidence. In fact, inspecting Figure 3B it is possible to see participants who were significantly less accurate on high PE trials yet were significantly more confident (upper left quadrant), indicating a dissociation between accuracy and confidence. Although our main goal was to equate accuracy across different confidence levels, which was successful, we also examined whether the PE manipulation led to changes in RT. As show in in Figure 3B, most participants were also faster for high PE (mean RT = 0.97±0.04s) compared to low PE (mean RT = 1.05±0.05s), leading to a significant difference (*t*(24) = 4.32, *p* < .001). Moreover, individual differences in the PE effect on confidence were negatively correlated with the PE effect on RT (*rho* = -.55, *p* = 0.005). Thus, even when dissociating confidence from accuracy, there is still a close link between confidence and RT, as is occasionally observed using this manipulation (Samaha et al., 2016).

We next turned to the neural data, which was analyzed in the same way and using the same central parietal electrodes as in Experiment 1. We first sought to replicate Experiment 1 and confirm that the CPP behaved as expected for an evidence accumulation signal. As shown in Figure 4A, the CPP slope was inversely correlated with RT such that a longer RT was associated with a shallower CPP spanning a significant cluster from 260 to 400 ms relative to stimulus onset and -510 to -100 relative to response onset. The CCP slope also predicted accuracy such that the CPP ramped up faster for correct compared to incorrect trials between 300 and 450 ms post-stimulus and between -720 and -680 and from -250 and -200 prior to the response (Figure 4B). Additionally, the CPP also predicted confidence ratings, with steeper slopes on high compared to low confidence trials (mean-split) from 360 to 420 ms post-stimulus and -500 to -180 ms pre-response (Figure 4C). Critically, we observed a similar effect when comparing high and low PE trials, which were matched for accuracy, but which differed significantly in confidence. In particular, high PE trials were associated with steeper CPP slopes than low PE trials in a significant cluster spanning 180 to 220 ms and 300 to 450 ms post-stimulus and -490 to -420 ms and -250 to -200 ms pre-response (Figure 4D). These findings collectively replicate and extend Experiment 1 by showing that confidence is correlated with pre-decision evidence accumulation signals even when accuracy is statistically equated.

**Figure 4.**
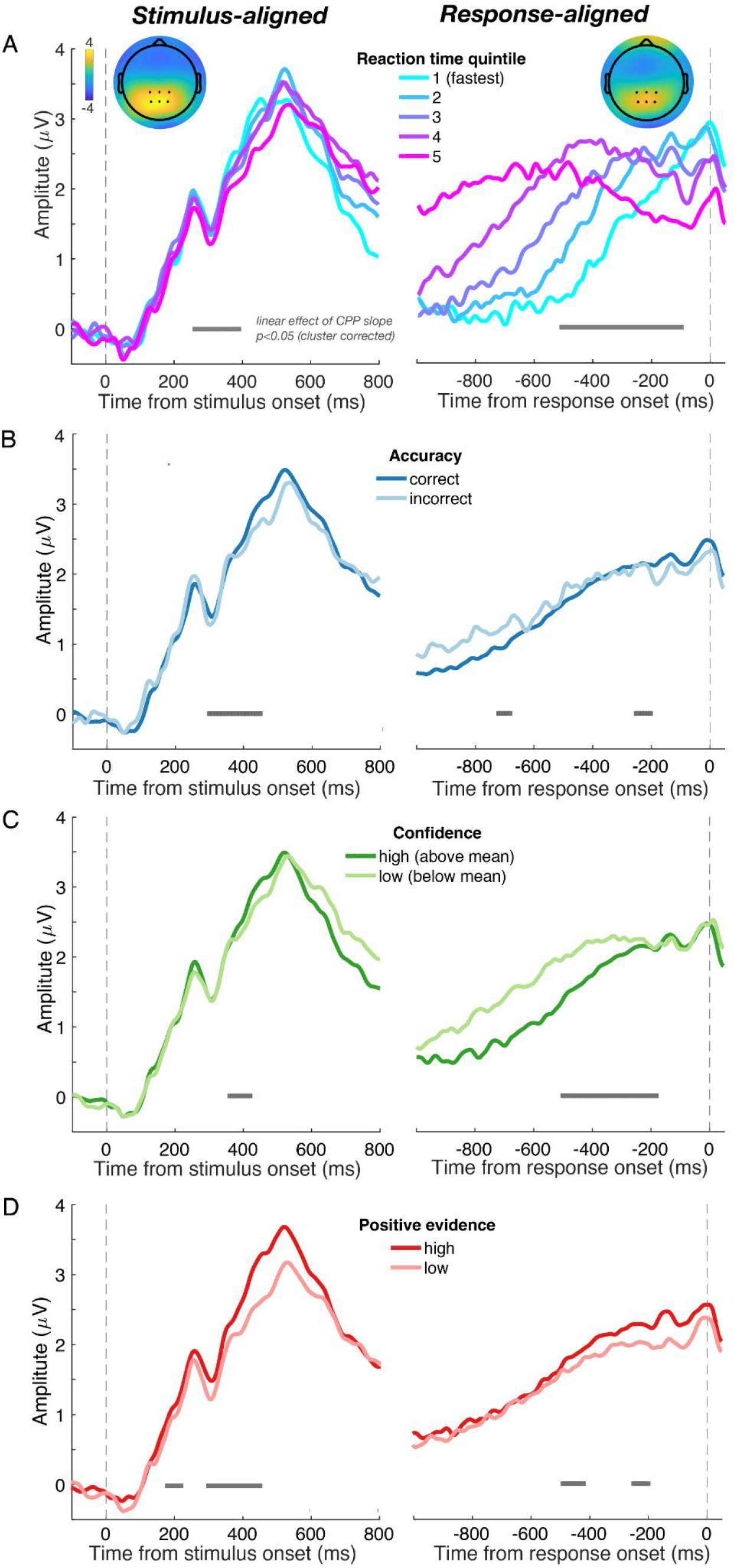
Grand average ERP time-courses aligned to both the target onset (left column) and response execution (right column) for Experiment 2 (*n* = 25), where the positive evidence bias (PEB) was introduced. (A) Both stimulus-aligned and response-aligned CPP slopes increased with faster RTs (5 equally spaced bins). (B) Both stimulus-aligned and response-aligned CPP slopes were higher on correct trials than on incorrect trials. (C) Both stimulus-aligned and response-aligned CPP slopes were steeper on high confidence (above mean) trials than on low confidence (below mean) trials. (D) Both stimulus-aligned and response-aligned CPP built up faster on high PE trials than on low PE trials - two conditions that did not differ in accuracy (see Figure 3).

### Statistical isolation of confidence signals from decision RT

Experiments 1 and 2 leave open the possibility that CPP reflects RT-related signals rather than confidence since the PE manipulation successfully controlled for accuracy only. In those experiments, however, participants indicated their direction decision and confidence using a single button press (see Figure 5A). This choice was intentional in order to increase the chances that the decision and confidence response were based on the same accumulated evidence (i.e., participants could not make their choice then continue accumulating evidence, from memory for instance, before rating confidence). However the single-response design means that RTs reflect a compound measure of decision+confidence time and are difficult to interpret. Indeed, inspecting the response-aligned data from Experiments 1 and 2 (Figures 2 and 4), evidence accumulation does not seem to always increase monotonically prior to the response but sometimes dips down before increasing again just prior to the button press (perhaps reflecting a sequential direction then confidence decision). To investigate if the CPP predicts confidence while statistically controlling for RT, we required a cleaner measure of decision RT to time-lock our data to and to use in a multiple regression model.

**Figure 5.**
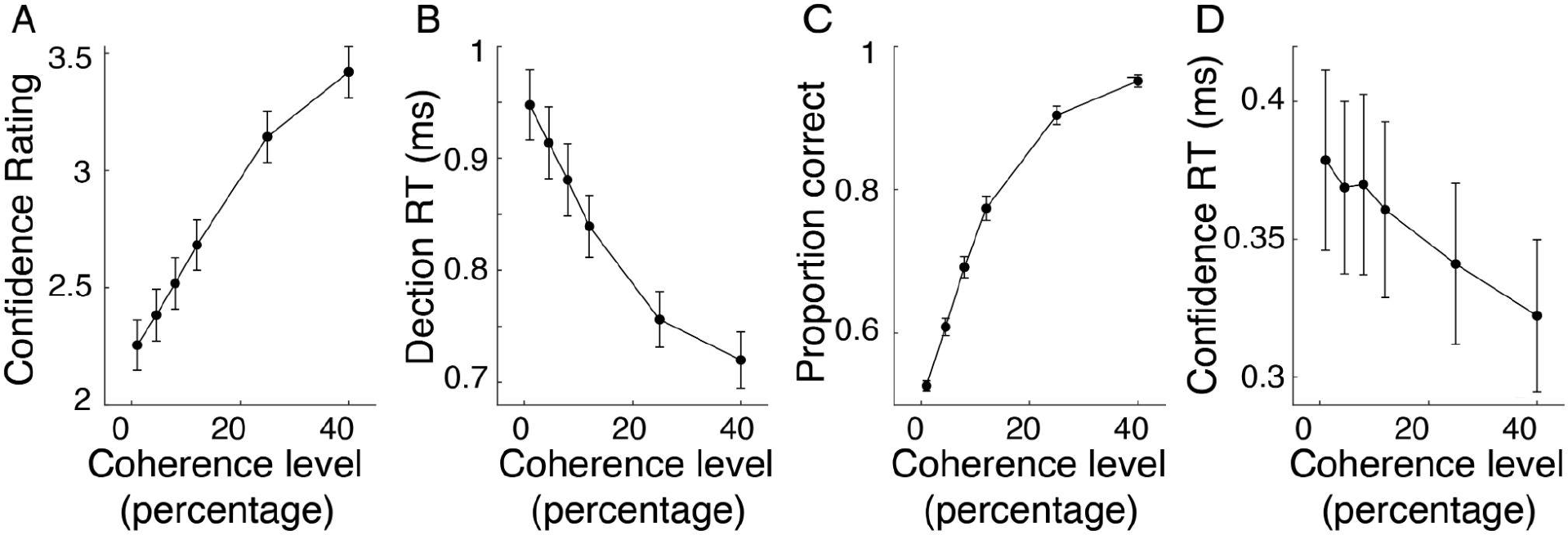
Behavioral results for Experiment 3 (*n* = 24). Confidence rating (A) and proportion correct (C) increased with higher coherence level. Decision RT (B) and confidence RT (D) were faster at higher coherence levels. Error bars denote ±1 SEM (across subjects)

In Experiment 3, participants (*N* = 24) reported motion direction and confidence sequentially while coherence was manipulated. This gives us a decision RT measure to use in a statistical model factoring out unique variance in confidence explained by CPP slope. First, the behavioral data showed that motion coherence had a significant effect on accuracy (*F*(5,115) = 409, *p <* .001), decision RT (*F*(5,115) = 80.85, *p <* .001), confidence (*F*(5,115) = 66.19, *p <* .001), and confidence RT (*F*(5,115) = 7.37, *p <* .001; See Figure 5B), as expected.

We then conducted the same ERP analysis as in Experiments 1 and 2 to confirm that the CPP predicted behavior in the expected way using a cleaner RT measure. We found that coherence level significantly modulated the build-up rate of the CPP in a cluster spanning 200 to 470 ms post-stimulus and -500 to -130 ms prior to response execution (Figure 6A). Decision RT varied with CPP slope in a cluster from 200 to 440 ms for stimulus-aligned CPP and a cluster from -500 to -150 ms for response-aligned CPP (Figure 6B). The CPP on correct trials ramped up faster than it did on incorrect trials within a cluster from 280 to 460 ms for stimulus-aligned data and -480 to -160 ms for response-aligned data (Figure 6C). Lastly, we also replicated the finding that CPP slope rate was significantly steeper on high compared to low confidence trials from 280 to 460 ms post-stimulus onset and from -510 to -170 ms pre-response (Figure 6D). Thus, with a more precise decision RT estimate we found that pre-choice evidence accumulation signals predict perceptual confidence. Moreover, the CPP more clearly followed a monotonic rise compared to Experiments 1 and 2, as expected of an evidence accumulation signal.

**Figure 6.**
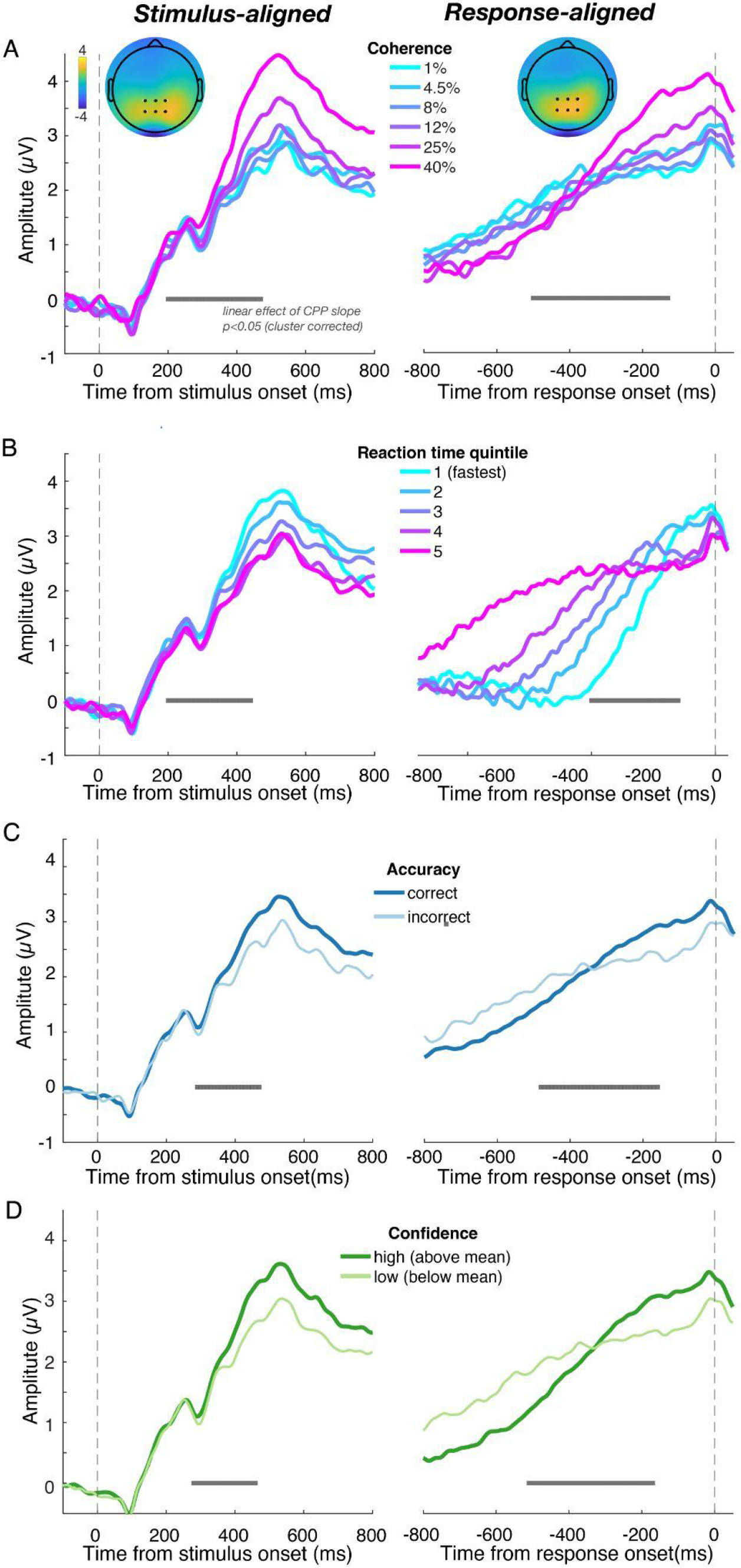
Grand average ERP time-courses aligned to both the target onset (left column) and response execution (right column) for Experiment 3 (*n* = 24), where choice and confidence responses were made sequentially. The results replicated what we found in Experiment 1 and in Experiment 2 showing that slopes of stimulus-aligned and response-aligned CPP both scaled with (A) coherence level, (B) decision RT, (C) accuracy, and (D) confidence.

To attempt to statistically control for the influence of coherence, accuracy, and RT on the relationship between CPP slope and confidence, we turned to a single-trial regression framework for our EEG data (Cohen & Cavanagh, 2011). For each trial, we quantified the CPP slope as a line fit to the EEG signal between 250 and 500 ms relative to stimulus onset and -300 to -100 ms relative to the response. Then, for each participant, we fit a general linear model predicting the single-trial CPP slopes using motion coherence, accuracy, RT, and confidence as predictors in the same model. We then extracted the t-statistic for each predictor and participant and performed a t-test at the group level to assess whether each predictor was reliably different from zero across the sample of participants. By using all predictors in the same model, we can statistically control for the effect of RT, coherence, and accuracy on CPP slope and identify the unique effect of confidence. However, any null effects should be interpreted cautiously since correlation between the predictor variables can lead to an inflation of the predictor variances and underestimation of effect sizes of the regression estimates, making it more difficult to observe a significant effect (i.e., collinearity inflates type-2 errors but not type-1 errors; Lavery et al., 2019)

Using stimulus-aligned single-trial slope estimates, the model showed a significant effect of coherence (*t*(23) = 7.46, *p* < .001) and a significant effect of confidence (*t*(23) = 2.15, *p* = .04) on slope. For response-aligned data, coherence level (*t*(23) = 2.57, *p* = .02), confidence (*t*(23) = 2.48, *p* = .02), and RT (*t*(23) = -3.33, *p* < .01) were all significant predictors (see Figure 7). The results indicate that higher confidence trials are associated with faster build-up of single-trial EEG signals, even when stimulus strength, RT, and discrimination accuracy are jointly modeled. In other words, confidence predicts unique variation in the neural signatures of evidence accumulation, above and beyond that explained by RT, accuracy, and stimulus evidence.

**Figure 7.**
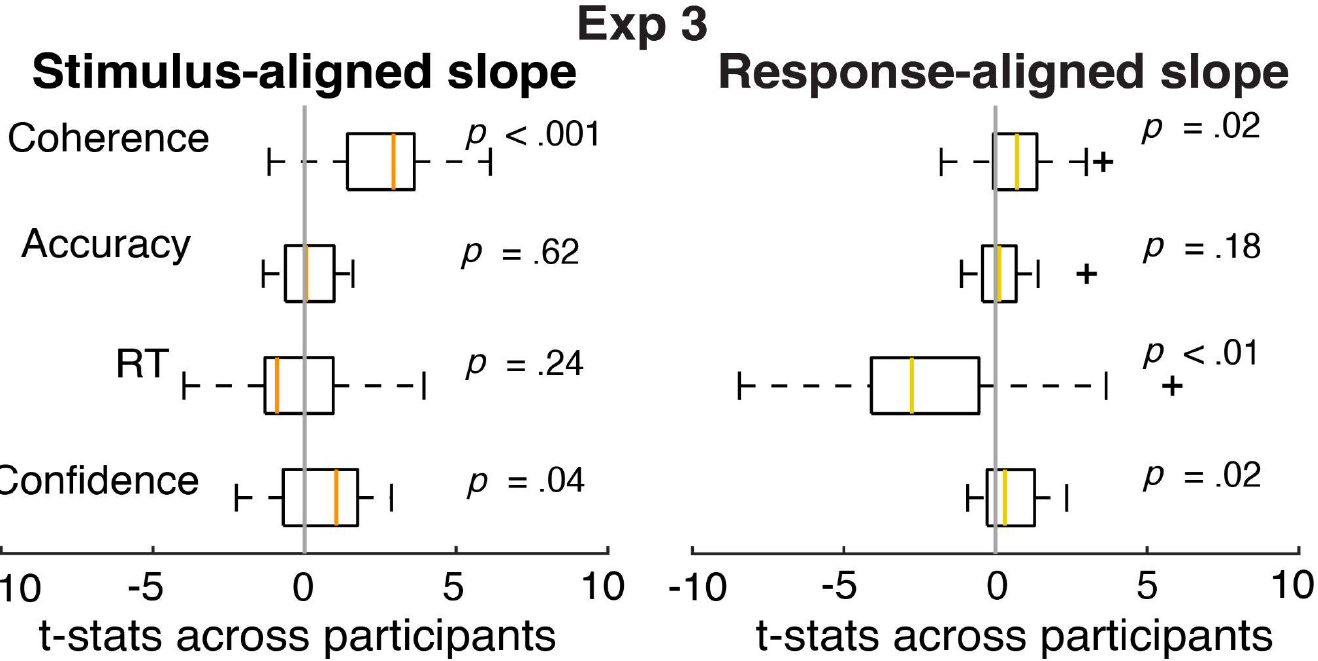
Results of a single-trial multiple regression analysis of EEG data from Experiment 3 (*n* = 24). Boxplots showing the distribution of t-statistics across participants for each predictor (e.g. coherence, accuracy, RT, and confidence) from a general linear model predicting the slopes of stimulus-aligned single-trial EEG (within the time window of 250 ms to 500 ms; left plot) and response-aligned single-trial EEG (within the time window of -300 ms to -100 ms; right plot). Orange and tan lines represent the median t-stat across participants for stimulus- and response-aligned analyses, respectively and for each predictor. The box boundaries represent the 25th and 75th percentiles of the t-statistics across participants and the whiskers represent the max and min value of the t-stat without an outlier (outliers identified with a “+”). Coherence and confidence predicted stimulus-aligned EEG slope controlling for other predictors in the model and coherence, RT, and confidence predicted response-aligned EEG slope, controlling for other predictors.

## Discussion

To characterize how confidence arises in perceptual decision making, we tested the hypothesis that confidence reflects the state of accumulated evidence leading up to a perceptual choice.

Across three experiments, we demonstrate that neural signatures of evidence accumulation, as reflected in the CPP, predict decision accuracy, RT, evidence strength, and subjective confidence judgments. There is currently little consensus regarding the neural computations underlying confidence judgments with some models linking confidence to objective evidence quality, and thus accuracy (Geurts et al., 2022; Meyniel et al., 2015; Sanders et al., 2016), and other models postulating a mapping from evidence and decision time to confidence (Fetsch et al., 2014; Gherman & Philiastides, 2015; Herding et al., 2019b; Kiani et al., 2014; Kiani & Shadlen, 2009; Rausch et al., 2020; Tagliabue et al., 2019; Zylberberg et al., 2016). Our results, however, suggest the picture may be somewhat more complex. First, in Experiment 2, we found that confidence could be dissociated from accuracy and that, under such conditions, the CPP still built-up faster when subjective confidence was higher. This indicates that confidence computations are unlikely to be a faithful readout of the same evidence controlling decision difficulty. Secondly, in Experiment 3, we used a statistical model to show that variation in confidence was related to parietal evidence accumulation signals even when controlling for the effect of RT (and accuracy and evidence quality). Thus, RT information alone did not fully explain the link between confidence and the CPP.

What model could account for our pattern of results? In a series of experiments in humans and non-human primates, Zylberberg and colleagues introduced a doubly-stochastic RDM stimulus (Zylberberg et al., 2016). In addition to the typical random motion component of the RDM, the authors added differing amounts of random variation in the time-varying motion evidence itself, giving rise to motion stimuli with high and low “motion volatility”. Behaviorally, increasing volatility led to increased confidence and faster RTs, but no change in accuracy, much like the PEB effect we found here. Evidence accumulation models of their data suggested that increasing motion volatility increases variability of the within-trial evidence accumulation process such that the decision variable reaches a choice boundary more rapidly, giving rise to the faster RTs observed in behavior. However, since increasing evidence volatility does not change the mean evidence strength, the decision variable was no more or less likely to hit the correct choice boundary. As a consequence, accuracy was not strongly impacted. Finally, to explain the increase in confidence associated with higher evidence volatility, the authors fit a model whereby the mapping from accumulated evidence to confidence was not adjusted appropriately to account for the change in volatility. In other words, observers seemed to apply a fixed mapping between decision time and confidence, one that did not take into account within-trial variability in the accumulation process. This account seems well-suited to explain the results from our Experiment 2, where changes in confidence absent changes in accuracy were associated with more rapid evidence accumulation signals in the brain. The implication of this is that increasing PE and NE effectively changes within-trial evidence accumulation variability such that a choice boundary is reached sooner and RTs are faster. Decision-makers then apply a learned mapping from decision time to confidence that is no longer entirely appropriate and they, essentially, assume that the faster RT was due to higher-quality evidence. Although many studies have used the PEB to study confidence behavior (Koizumi et al., 2015; Maniscalco et al., 2016; Rollwage et al., 2020; Samaha et al., 2016, 2019; Samaha & Denison, 2022; Zylberberg et al., 2012), or build neurally-plausible models of confidence (Khalvati et al., 2021; Maniscalco et al., 2021) our results reveal a novel neural correlate of the PEB in perceptual confidence.

The above-mentioned framework, however, appears to be inadequate to explain the results of our Experiment 3, wherein we found that residual variance explained in CPP slope was related to confidence even when controlling for accuracy and RT. This finding suggests that confidence is obviously related to accuracy and RT, but still partially separable from both of them and that confidence computations could be more complicated than just mapping the amount of accumulated evidence and how long it took to accumulate that evidence onto a subjective estimate of confidence. Evidence accumulation models typically assume that a certain proportion of the response time is not accounted for by the decision process, but is instead related to sensory-encoding and motor execution processes, this is referred to as non-decision time. The measured RT, then, is the sum of the non-decision time and the decision time. Therefore, one possibility is that trial-to-trial variability in non-decision time (sensory encoding latencies, for instance) contributes to RT but not to the evidence accumulation rate on that trial. As a result, there will be partial independence between RT and evidence accumulation rate such that controlling for RT does not necessarily fully control for decision time. The best approach for modeling non-decision time (e.g., as fixed or variable across trials) remains an issue of debate (Bompas et al., 2023; Weindel et al., 2021)

A second possibility to explain the persistent link between CPP slope and confidence after controlling for RT is that post-decisional factors influence confidence as well. Many recent models propose that evidence continues to accumulate after a decision is made and that confidence is determined by the amount of choice-consistent or choice-inconsistent post-decision evidence (Desender et al., 2019; Desender, Ridderinkhof, et al., 2021; Navajas et al., 2016; Pleskac & Busemeyer, 2010; Yu et al., 2015). Although our results clearly indicate that subjective confidence can be predicted by pre-decision evidence signals, we cannot rule out that confidence is also informed by post-decisional dynamics. We did not systematically investigate post-decision signals in our study because a cursory view of the response-locked ERPs indicates that the CPP appears to decline immediately after the response, rather than continue accumulating. This is likely because we used a fixed-duration stimulus of 300ms in all studies, whereas many experiments looking at post-decision evidence either continue to present evidence after the choice or right up until the choice. There is also some uncertainty regarding the correct baseline procedure to use. If response-locked ERPs are baselined relative to a pre-response window, then any differences leading up to the response (which we observe) will trivially appear in the post-decision window, mimicking post-decision evidence signals (Feuerriegel et al., 2022). Nevertheless, our data suggest that pre-decision evidence accumulation signals alone can predict confidence even when controlling for accuracy, RT, and evidence strength, however there may still be a contribution of post-decision signals to the final confidence judgment.

## Methods

### Participants

25 participants (twelve men; age range 18-30) completed Experiment 1. 26 participants (six men; age range 18-40) completed Experiment 2 (one was removed from analyses due to using the wrong response keys in the final blocks of the task). 25 participants (eight men; age range 18-31) completed Experiment 3 (one was excluded due to rating a confidence level of 4 on 96% of trials). All participants were recruited from the University of California, Santa Cruz (UCSC) for course credit. They reported normal or corrected-to-normal vision and provided written informed consent. All procedures were approved by the institutional review board at UCSC.

### Stimulus and Apparatus

In all experiments, stimuli were presented on a black background using an electrically-shielded VIEWPixx/EEG monitor (120 Hz refresh rate, 1920 × 1080 pixels resolution) that was ∼53 cm wide and viewed at a distance of ∼69.5 cm from a chinrest. Stimulus presentation and behavioral data were controlled by Psychtoolbox-3 (Kleiner et al., 2007; Pelli, 1997) running in the MATLAB environment.

In all experiments, the target stimulus consisted of 150 white dots (0.03 degrees each) presented within a 5 degree aperture centered on fixation. For each stimulus, a percentage of the dots (referred to as the motion coherence level) was randomly selected on each frame to be displaced by a fixed distance of 0.042 degrees (corresponding to a motion speed of 5 degrees/second) in either the left or right direction on the following frame. The rest of the dots were placed randomly and independently within the circular aperture. A red dot (0.25 degrees of visual angle) on which participants were asked to fixate was superimposed atop the center of the dot stimuli. To prevent tracking of any single dot and encourage responses based on global motion direction, each dot had a lifetime of 66 ms.

In Experiments 1 and 3, motion coherence was chosen from one of the following 6 levels: 1%, 4.5%, 8%, 12%, 25%, or 40% with equal probability. In Experiment 2, we created high and low positive evidence (PE) stimuli by setting the coherence to be 50% in the high PE and 25% in the low PE conditions. We then used a 1-up, 3-down staircase procedure prior to the main task that varied the amount of negative evidence (NE, i.e., motion signal in the opposite direction as the PE) such that the high and low PE trials both approximated ∼79% accuracy.

### Procedures

*Experiment 1.* Participants were tested individually within a dim, sound attenuated room. The experimenter explained the instruction to the participant and verified that the participant understood the instructions before proceeding to the practice trials and critical blocks. The participant completed a practice block with 180 trials using an easier set of coherence levels than the critical task (5%, 30%, 40%, or 70%). On each practice trial, a beep sound was presented if participants incorrectly judged the motion direction. No feedback was presented on the critical trials. Participants proceeded to the critical blocks only if they achieved 80% accuracy or greater on the 70% coherence practice trials. Participants who failed to reach this criteria in the very first practice block performed more practice blocks until they met the criteria. This criteria was set to make sure that participants fully understood the task and could perceive the motion signals.

In critical trials, participants performed a random dot motion discrimination task (Roitman & Shadlen, 2002; Kelly & O’Connell, 2013) with confidence ratings. Participants were told that they would be presented with a group of moving dots on each trial, with some dots moving toward either the left or right and the rest moving randomly. Participants were instructed to report the global direction of motion along with their confidence as quickly and accurately as possible. Each trial began with a red fixation point presented in the center of the screen for a random inter-trial interval between 1000 and 1500 ms. The dot motion stimulus then appeared for 300 ms, with the red dot remaining on the screen until participants pressed a single key to indicate both the direction of the global motion and their confidence rating on a 4 point scale. We informed participants that “1” denotes a random guess at the motion direction and “4” denotes high confidence in their decision. Participants were instructed to keep the fingers of their left hand (from pinky finger to index finger, respectively) on keys ‘A’, ‘S’, ‘D’, and ‘F’ which represented left motion with the confidence rating from 4 to 1. They also kept the fingers of their right hand on keys ‘J’, ‘K’, ‘L’, and ‘;’, representing right motion with confidence from 1 to 4, respectively.

Each participant completed 1080 trials in total, consisting of 180 trials within each motion coherence level. The trials were presented in six blocks with 180 trials in each block. Motion direction and coherence level varied pseudo-randomly on a trial-by-trial basis.

*Experiment 2.* Participants first performed 80 trials of practice with the 50% and 25% coherence PE stimuli with low NE coherence levels (10% and 1.5%, respectively). Auditory feedback was given after each error during the practice. If overall performance was below 80% after the first practice block, a second was administered. Following practice, 200 trials of an adaptive staircase procedure were performed wherein the NE coherence increased by 3% following a correct response and decreased by 3% following three incorrect responses. Two separate staircases were interleaved for the high and low PE stimuli in order to find NE thresholds for each PE level that approximate 79% accuracy. Following the staircase, 1080 trials of the critical task were run split into 6 blocks. NE levels for high and low PE stimuli were fixed throughout the critical task and high or low PE stimuli were randomly presented on each trial. Participants provided responses in the same manner as Experiment 1.

*Experiment 3.* The procedure was identical to Experiment 1 except that participants were instructed to report the motion direction and confidence sequentially. After the stimulus disappeared, participants either pressed the ‘<’ key using the right index to indicate leftward motion or the ‘>’ using the right middle finger to indicate rightward motion. Participants then pressed one of the 1-4 number keys across the top row of the keyboard using their left hand in order to indicate their confidence. Participants completed 1080 trials split into 6 blocks.

### EEG recording and analysis

In all experiments, continuous EEG was acquired from 63 active electrodes (BrainVision actiCHamp), with impedance at each central-parietal electrode kept below 20kΩ. Recordings were digitized at 1000 Hz and FCz was used as the online reference. EEG was processed offline using custom scripts in MATLAB (version R2019b) and with EEGLAB toolbox (Delorme & Makeig, 2004). Data were high-pass filtered at 0.1 Hz and low-pass filtered at 30 Hz using a zero-phase Hamming-windowed sinc FIR filter and then downsampled to 500 Hz. Data were re-referenced offline to the average of all electrodes. Continuous signals were then segmented into epochs centered on stimulus onset using a time window of -2000 ms to 2000 ms. Individual trials were rejected if eye-blinks occurred during stimulus presentation or if any scalp channel exceeded ±100 μV at any time during the interval extending from -500 to 500 ms relative to the stimulus onset. On average, 209 trials were rejected for each participant in Experiment 1, 130 trials were rejected for each participant in Experiment 2, and 140 trials were rejected for each participant in Experiment 3. Trials excluded from the EEG data were also not involved in the analysis of behavior. Noisy channels were spherically interpolated and an independent components analysis (infomax algorithm) was performed to identify and subtract one or two components per subject that reflected eye blinks or eye movements. Lastly, a pre-stimulus baseline of -200 ms to 0 ms was subtracted from each trial.

Based on the inspection of the grand average scalp topographies from Experiment 1 (within the time window of 250 ms to 450 ms post-stimulus onset and -510 ms to -130 ms prior to response execution), we chose the following electrodes for all subsequent analysis across all experiments: CPz, CP1, CP2, Pz, P1, and P2. These electrodes captured the maximum CPP amplitude and were chosen based on the grand average waveform, independent of any subsequent comparison between conditions.

### Statistical analyses

In all experiments, the CPP build-up rate was defined as the slope of a line fit to each participant’s average CPP waveform (Twomey et al., 2015) using a 200 ms sliding window advanced in 10 ms steps between 100 ms to 550 ms relative to the stimulus-aligned CPP waveform and -1000 ms to -10 ms relative to the response-aligned CPP waveform. We then fit separate linear models predicting CPP slopes at each time window by motion coherence level, RT (divided into 5 quantiles), accuracy (correct or incorrect), confidence (high or low based on a participant-specific mean-split), and positive evidence (high or low; Experiment 2 only). For each model we compared the regression slopes at each time window to zero (one-tailed) and used a nonparametric, cluster-based permutation test to correct for multiple comparisons across time (Maris & Oostenveld, 2007). Specifically, we stored the largest cluster of significant timepoints across each of 10,000 permutations of the data and only considered clusters in the real data that surpassed the 95 percentile of this distribution of cluster sizes expected under the null hypothesis.

In Experiment 3 we also ran a single-trial multiple regression model (Cohen & Cavanagh, 2011) by fitting a line to single-trial EEG signal between 250 and 500 ms relative to stimulus onset and from -300 to -100 ms relative to the response. We stored the beta coefficients (slopes) for each trial and then fit a linear model predicting single-trial EEG slopes using confidence, coherence level, accuracy, and RT as the predictors. Single-trial slopes and RTs were both rank-scored in the model to mitigate the effect of potential outliers while still testing for monotonic relationships. Thus, the model had the form Y = INT + *b_1_*CONF + *b_2_*COH + *b_3_*RT + *b_4_*ACC where Y is rank-scored single-trial EEG build-up rates, INT is the intercept, *b_1–4_*are regression coefficients, and CONF, COH, RT and ACC are trial vectors of the participant’s confidence, coherence level, rank-scored reaction time, and accuracy on each trial. The resulting t-value of each predictor for each participant indicates the normalized effect of that predictor on single-trial EEG slopes. These t-values were saved for each participant and predictor and tested against zero at the group level using two-tailed, repeated-measures t-tests.

## Notes

### Competing Interest Statement

The authors have declared no competing interest.

